# Drivers of thermal tolerance breadth of plants across contrasting biomes

**DOI:** 10.1101/2023.10.02.560437

**Authors:** Verónica F. Briceño, Pieter A. Arnold, Alicia M. Cook, Stephanie K. Courtney Jones, Rachael V. Gallagher, Kris French, León A. Bravo, Adrienne B. Nicotra, Andy Leigh

## Abstract

1. The climate variability hypothesis (CVH) predicts that species from environments with more variable temperatures should have wider thermal tolerance breadth. This hypothesis has not yet been tested thoroughly across diverse plants. Here, we asked how local climate predictors (including precipitation, mean and extreme temperatures and thermal variability) are associated with species physiological thermal limits.
2. Measures of lower (T_crit-cold_) and upper (T_crit-hot_) photosystem II thermal tolerance thresholds were used to determine thermal tolerance breadth (TTB), along with ice nucleation temperature (T_nucleation_, freezing tolerance) of 69 plant species sampled from the field across three contrasting biomes: alpine, desert and coastal temperate rainforest.
3. All measured thermal tolerance metrics (T_crit-cold_, T_nucleation_, T_crit-hot_ and TTB) differed among biomes. Notably, desert species had the most cold and heat tolerant leaves, and therefore the widest TTB, whereas species in alpine and temperate biomes had similar TTB. For plants in all biomes, TTB exceeded the thermal range of their local climate.
4. Overall, two Principal Component axes of local climate drivers explained substantial variation in all tolerance metrics. Extreme hot, dry climates improved freezing and heat tolerance. High thermal variability and low minimum temperatures also improved freezing tolerance but were unrelated to heat tolerance or TTB. Species explained a significant amount of variation among all metrics, but this was not due to phylogenetic relatedness. We discuss how the remaining variation could be due to microclimate-driven plasticity, leaf traits or thermoregulatory mechanisms.
5. **Synthesis**. Our results provide partial support for the climate variability hypothesis in plants: photosystem thermal tolerance breadth was greatest in more thermally variable biomes. This relationship was largely driven by cold tolerance, with variation in heat tolerance explained better by mean and extreme temperatures. Therefore, we conclude that the CVH alone is not sufficient to explain variation in plant thermal tolerance, with many other aspects of climate, environment and biology being potentially important drivers.

## Introduction

Desert and alpine plant species live close to the thermal limits of biological processes. Unlike those in temperate regions, plants living in extreme environments are exposed to both very low and very high temperatures. Given the important influence of temperature in shaping species distributions (Tattersall *et al*. 2012; Garcia-Pichel *et al*. 2013), the innate thermal tolerance thresholds (the lowest and highest temperature that a species can withstand) and thermal tolerance breadth (TTB; the range of temperature between upper and lower thermal tolerance thresholds) of species might reflect the temperature variation that characterises their environment. The climate variability hypothesis (CVH; Janzen 1967) predicts that species with relatively wide thermal tolerance ranges are likely to evolve in environments with variable temperatures compared to thermal specialists with relatively narrow tolerance ranges that evolved in thermally stable environments.

The CVH has been applied across many environmental gradients (e.g. elevation, latitude, climate) and timescales of thermal variability (e.g. Gutiérrez-Pesquera *et al*. 2016; Shah *et al*. 2021; Bovo *et al*. 2023), thus it can be interpreted differently. Here, we follow the interpretation that the *annual temperature range* has shaped thermal tolerance breadth over evolutionary time, allowing organisms to tolerate *seasonal shifts* in temperatures (Kefford *et al*. 2022; Dewenter *et al*. 2024). The CVH is a cornerstone of thermal ecology; it has been tested broadly on animals (Addo-Bediako, Chown & Gaston 2000; Compton *et al*. 2007; Calosi *et al*. 2010; Baudier *et al*. 2018) but on relatively few plant species. There is also limited consistent evidence supporting the CVH in plants, and even studies on the same genus show mixed support (e.g. *Mimulus*; Wooliver, Tittes & Sheth 2020; Querns *et al*. 2022; Chiono & Paul 2023).

Cold and heat tolerance thresholds that define the bounds of TTB are rarely studied concomitantly in plants (Geange *et al*. 2021). In natural systems, cold tolerance studies are dominant in alpine environments where plants can withstand very low temperatures and tolerate extracellular ice formation and the resulting cytoplasmic dehydration (Sakai & Larcher 1987; Larcher 2003). For example, alpine plants are known to tolerate extreme cold, many avoiding damage below –5 °C or –10 °C (Neuner & Pramsohler 2006; Harris *et al*. 2024) and resisting freezing below –15 °C (Venn, Morgan & Lord 2013; Neuner *et al*. 2020) or even below –30 °C (Bannister *et al*. 2005). Heat tolerance studies have historically been primarily focussed on warm and desert environments, where heat tolerance has often been associated with native microhabitat conditions (Knight & Ackerly 2002; Curtis *et al*. 2016). Desert plants exhibit tolerance to heat often in excess of 50 °C (Monson & Williams 1982; Downton, Berry & Seemann 1984; Seemann, Berry & Downton 1984; Zhu *et al*. 2018), and some species have a remarkable ability to tolerate over 55 °C during heatwave conditions (Andrew *et al*. 2023). Temperate plants tend to have less capacity to tolerate heat and cold, many only tolerating cold to –5 °C (Bannister & Lord 2006) and heat to 45 °C (Bilger, Schreiber & Lange 1984; O’Sullivan *et al*. 2017). Despite focus on either cold or heat tolerance in either climate extreme, plants in these biomes can be exposed to both cold and heat extremes, even within growing seasons. As the climate continues to warm and extreme climatic events increase, understanding the cold and heat tolerance of plants that grow in a range of biomes will be essential (Geange *et al*. 2021).

Temperature increases due to climate change has motivated an increase in comparative studies and syntheses of plant thermal tolerance, particularly in relation to climate variability and the occurrence of increasing extreme climatic events (Rummukainen 2012; Seneviratne *et al*. 2021; Andrew *et al*. 2023; Harris *et al*. 2024). An assessment of plant heat tolerance in four continents showed a strong pattern of heat tolerance increasing as the mean maximum temperature of the warmest month increased (O’Sullivan *et al*. 2017). A global meta-analysis encompassing a wide range of species and biomes suggests that cold tolerance increases towards the poles and heat tolerance increases towards the equator, where the latitudinal cline is a coarse proxy for mean annual temperature (Lancaster & Humphreys 2020). However, substantial variation in these tolerance values is due to methodological differences among studies and locations (Perez *et al*. 2021). Heat and cold tolerance are rarely measured together within the same study, let alone with equivalent methods, but both these aspects are important for accurately and precisely quantifying thermal tolerance breadth (Geange *et al*. 2021; Perez *et al*. 2021).

Differing patterns of climate among contrasting biomes provide an opportunity to test the CVH by exploring the role of climate variability in upper and lower thermal tolerance thresholds and on thermal tolerance breadth. While broad analyses indicate that maximum temperatures can influence heat tolerance (O’Sullivan *et al*. 2017) and mean annual temperature influences both cold and heat tolerance (Lancaster & Humphreys 2020), the climate predictors of thermal tolerance breadth, are virtually unknown for plants. Given that water availability affects both plant heat tolerance (Curtis *et al*. 2016; Cook *et al*. 2021; Marchin *et al*. 2022) and freezing tolerance (Sierra-Almeida, Reyes-Bahamonde & Cavieres 2016), precipitation is also likely to influence thermal tolerance breadth. If we can reveal general patterns underlying variation in, and the climate drivers of, thermal tolerance thresholds and breadth among co-occurring plant species within and among biomes, we can strengthen predictions of which species and where will likely be most vulnerable to changing thermal conditions.

Here we sought to determine the climate drivers of leaf-level physiological thermal tolerance thresholds and thermal tolerance breadth of plant species growing in extreme and comparatively benign environments in the context of the climate variability hypothesis (CVH). Our field campaigns covered three biomes found in south-eastern Australia that contrast strongly in their thermal environments. For simplification, hereafter we refer to these as alpine, desert and temperate biomes. The alpine biome is characterised by both strong daily temperature oscillations and strong temperature seasonality, with a cold winter and relatively mild summer; the desert biome is arid, has strong daily temperature oscillations and strong temperature seasonality, with particularly hot summers; the temperate biome has relatively mild daily and seasonal oscillations, with mild winters and high humidity year-round (Loidi, Navarro-Sánchez & Vynokurov 2022).

We used chlorophyll fluorescence to determine the low and high photosystem II (PSII) thermal tolerance thresholds and TTB of 22 alpine, 24 desert and 23 temperate species during the growing season. To examine the relationships between climate variability and thermal tolerance thresholds, we extracted long-term, locally downscaled climate parameters for each sampling locality. We addressed three objectives in this study. First, we assessed if and how the thermal tolerance thresholds and TTB of a broad range of plant species differed among the contrasting biomes. We hypothesised that thermal tolerance thresholds would reflect the climate of the respective biomes. That is, colder threshold temperatures in the alpine and hotter threshold temperatures in the desert, with temperate species having both milder cold and heat tolerance thresholds. Second, we tested whether species from the three biomes differed in their TTB. We hypothesised that species from the more thermally variable biomes (alpine and desert) would have wider thermal tolerance breadth than those from the thermally stable biome (temperate), in accordance with the CVH. Third, we assessed whether variation in thermal tolerance thresholds and TTB could be explained by local climate variables, and whether these relationships were linear or not. We hypothesised that climate extremes and variability would have a stronger association with thermal tolerance thresholds and TTB than mean annual climate values, supporting the CVH.

## Materials and Methods

### Study locations

Sites used in this study were designated to one of three biomes in New South Wales (NSW), Australia: alpine (Monaro Ngarigo, Ngunnawal, Wiradjuri and Walgalu Country, Kosciuszko National Park), desert (Ngemba and Barkindji Country, Gundabooka National Park) and temperate (coastal temperate rainforests of Dharawal, Wadi Wadi and Yuin Country in the Illawarra Region: Royal National Park, Illawarra State Conservation Area and Bass Point Conservation Area). This study was conducted under approval from the NSW Department of Planning, Industry and Environment (Scientific Licence SL102330). Sampling sites incorporated a range of vegetation types sampled in each biome (mean climatic details from sites are inset in Fig. 1). Our specific study biomes represent distinct domains, ecozones, biomes, and sub-biomes of the world by climatic definitions (Loidi, Navarro-Sánchez & Vynokurov 2022). Specifically, alpine falls within the cryocratic domain (sub-biome 1b: tundras of the temperate mountains in cryoro belt); desert is in the xerocratic domain (sub-biome 7b: temperate desert and semi-desert); and temperate is in the mesocratic domain (sub-biome 4a: lauroid evergreen forest of the lowlands) (Loidi, Navarro-Sánchez & Vynokurov 2022).

**Fig. 1.**
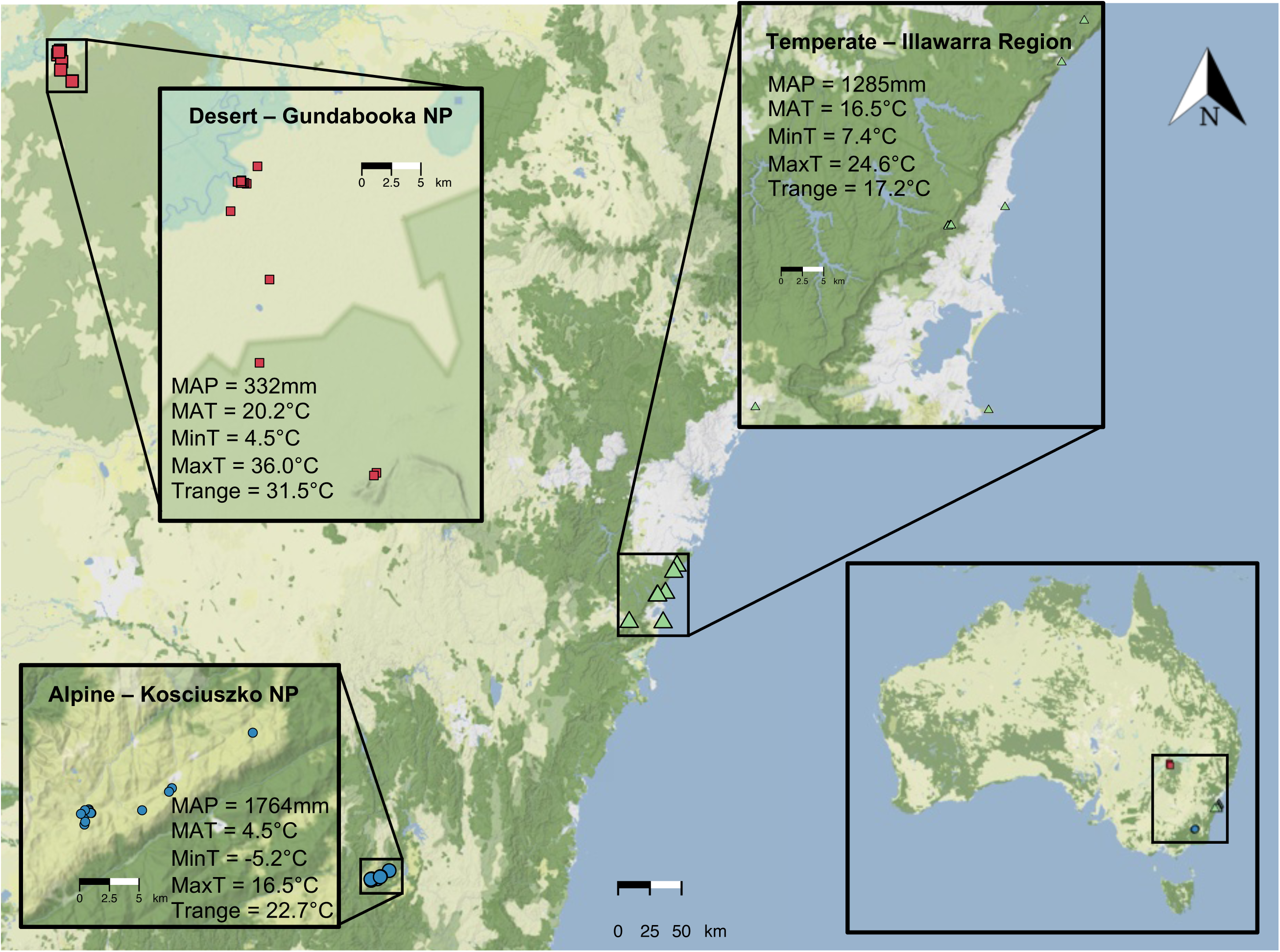
Locations of study sites across three biomes in New South Wales, Australia. Desert sites (red) in Gundabooka National Park (GNP), coastal temperate rainforest sites (green) in the Illawarra Region (temperate), and alpine sites (blue) in Kosciuszko National Park (KNP). GNP sites spanned open eucalypt woodland, mulga and chenopod shrublands between elevations of 100 to 224 m a.s.l. KNP sites encompassed feldmark, herbfields and sphagnum bog vegetation formations between 1792 and 1855 m a.s.l. In the temperate environments, collections were conducted across the Illawarra, including the tall open forest and rainforest communities between 100 to 400 m a.s.l. MAP = mean annual precipitation, MAT = mean annual temperature, MinT = mean minimum temperature of the coldest month, MaxT = mean maximum temperature of the warmest month, T_range_ = MaxT - MinT.

#### Species selection

A total of 69 species (22 alpine, 24 desert and 23 temperate) were selected to encompass a representative sample of growth forms and plant families and where possible, included congeneric and/or con-familial species between two or more biomes (Table S1). We included community dominants and a range of less abundant species to represent a cross section of species occupying each biome. Species replicates were sampled across days within each sampling period to incorporate daily variation in thermal tolerance thresholds.

#### Leaf sample collection

Mature, fully expanded leaves were harvested from five to seven individuals per species. For shrubs and trees, small branches were collected from the northern-facing side of the plant at a height of 1-1.5 m. For smaller growth forms, entire individuals or rosettes were collected. Samples were kept in cool sealed plastic bags with moistened paper towel until they were processed (approximately 1-2 h after collection). In the laboratory, leaves (size ranging from 0.25 cm^2^ to 5 cm^2^) were cut to around 1 cm^2^ of uniform photosynthetic tissue, avoiding prominent venation, damaged or discoloured areas or any structures that would reduce thermal contact of the leaf material during the thermal tolerance assays. For species with smaller leaves, we placed several leaves directly next to each other to fit an approximate area of 1 cm^2^. Leaf samples were kept on wet florist foam in humid dark containers until thermal tolerance assays were conducted. We selected when to collect field samples to capture the peak growing season in each biome; thus, sampling and thermal tolerance measurements occurred in January 2020 for the desert biome, March 2020 for the temperate biome, and December 2020 for the alpine biome.

### Thermal tolerance threshold assays

Photosystem II (PSII) is a protein complex embedded in the thylakoid membrane of plant cells that captures light energy during photosynthesis. PSII has long been recognised as a thermally sensitive structure (Schreiber & Berry 1977; Berry & Bjorkman 1980; Seemann, Berry & Downton 1984). Chlorophyll *a* fluorescence is light that is re-emitted by chlorophyll molecules in plants after absorbing light energy during photosynthesis, which provides valuable information about the efficiency and functioning of the photosynthetic process, especially during stress (Murchie & Lawson 2013). The temperature at which PSII is disrupted causes a rise in minimal chlorophyll *a* fluorescence (F_0_) as a leaf is heated or cooled (Schreiber & Berry 1977; Neuner & Pramsohler 2006). This critical temperature indicates a threshold beyond which physiological and photochemical systems become increasingly impaired, and damage can occur if temperatures are sustained.

#### Lower and upper thresholds of PSII (T_crit-cold_ and T_crit-hot_)

The critical lower and upper temperature thresholds of PSII (T_crit-cold_ and T_crit-hot_) were measured using the temperature-dependent (T) change in F_0_ (T-F_0_ curve) method, that is, the rise in F_0_ with set heating and cooling rates (Neuner & Pramsohler 2006; Arnold *et al*. 2021). One leaf per individual wherever possible (or multiple small leaves abutting one another for plants with very small leaves) was used for each of the heating and cooling assays (*n* = 5-7 replicates per species). For each set of T-F_0_ measurements, we secured 45 samples with double-sided tape to paper that was then placed on a Peltier plate (CP-121HT; TE-Technology, Inc., Michigan, USA) that was set to either cool or heat samples. The Peltier plate was controlled using LabView-based control software (National Instruments, Texas, USA) and adapted code from TE-Technology, Inc. Double-glazed glass was placed on top of the leaf samples on the plate to avoid water condensation, particularly at freezing temperatures, and to compress samples to ensure maximum leaf contact with the Peltier plate for uniform thermal transfer (see Arnold *et al*. 2021 for more details).

Chlorophyll fluorescence was measured with a Pulse Amplitude Modulated (PAM) chlorophyll fluorescence imaging system (Maxi-Imaging-PAM; Heinz Walz GmbH, Effeltrich, Germany) mounted above the Peltier plate. Leaves were first dark-adapted, and the PSII photochemistry maximum quantum efficiency (F_V_/F_M_) was measured to check the starting function of leaves before another dark adaptation period of 15 min prior to the assay. During temperature ramping, F_0_ was measured every 20 s with a weak blue low frequency (1 Hz) pulse modulated measuring light (0.5 μmol photons m^-2^ s^-1^). Leaf temperatures were simultaneously measured with 40-gauge type-T thermocouples (OMEGA Engineering, Singapore) attached to the undersides of leaves and logged with a multi-channel datalogger (DT85 DataTaker; Lontek, Australia). For hot T-F_0_ measurements, samples were heated from ambient temperature (15-20 °C) to 60 °C at the rate of 30 °C h^-1^ (i.e. 0.5 °C min^-1^, over ∼ 1.5 h). For cold T-F_0_ measurements, the leaves were cooled down from ambient temperature to at least –20 °C, at the rate of 15 °C h^-1^ (i.e. 0.25 °C min^-1^, over ∼ 2.5-3 h). These heating and cooling rates are moderate rates of temperature change that are considered appropriate for interspecific comparisons (Arnold *et al*. 2021). The starting ambient temperature of each heating/cooling assay depended on the temperature of the laboratory space used at each field site, which was generally ∼15 °C for alpine and temperate, and ∼20 °C for desert biomes.

T_crit-cold_ and T_crit-hot_ were each determined as the point of transition between slow-rise and fast-rise phases of F_0_ values that occur during steady cooling or heating (Fig. S1; see also Arnold *et al*. 2021). These critical temperature points were derived from breakpoint regression models fitted to each T-F_0_ curve, using the *segmented* R package (Muggeo 2017) (R code available from https://github.com/pieterarnold/Tcrit-extraction). T_crit-cold_ and T_crit-hot_ were used to represent the lower and upper bounds of the PSII thermal tolerance breadth, TTB (TTB = T_crit-hot_ – T_crit-cold_).

#### Ice nucleation temperatures

Freezing tolerance is tied to the temperatures at which ice formation occurs, called the ice nucleation temperature (T_nucleation_). Measuring T_nucleation_ is required to understand whether temperature-dependent damage to PSII occurs in association with ice formation. During T_crit-cold_ assays, the temperature of each leaf sample was measured continuously. T_nucleation_ can be observed as a small rise in leaf temperature release of heat energy (an exothermic reaction) when there is a sudden state transition and ice crystals form (i.e. nucleation). By plotting leaf temperature over time as samples cool, we identified the temperature at which the exothermic reaction initiated, which is T_nucleation_ (Larcher 2003).

### Local climate variables for sampling locations

Climate data were obtained from gridded datasets at 1 km resolution based on spatial interpolations of long-term (1981–2010) conditions accessed from the CHELSA v2.1 database (Karger *et al*. 2017). We extracted the following local climate variables at each location that the leaves of each species were collected: mean annual precipitation (MAP), mean annual temperature (MAT), mean minimum temperature of the coldest month (MinT), mean maximum temperature of the warmest month (MaxT). We calculated annual thermal range as: T_range_ = MaxT – MinT. Table S2 includes local climate variables for the complete list of species and sampling locations.

### Statistical analyses

All analyses and data visualisations were conducted in the R Environment for Statistical Computing version 4.5.1 (R Core Team 2023). Phylogenetic relatedness can contribute to explaining variation in physiological traits. Therefore, prior to model fitting, we generated a plausible phylogenetic tree using the R package *V.phylomaker* (Jin & Qian 2022) and calculated the phylogenetic signal and statistical significance for the four response variables (T_crit-hot_, T_crit-cold_, T_nucleation_ and TTB) using the *phylosignal* package (Keck *et al*. 2016). Both these tests of phylogenetic signal and incorporating phylogenetic relatedness in models described below allowed us to account for the shared evolutionary history across the 69 species that contributed to thermal tolerance thresholds.

To assess whether the five key local climate variables (MAP, MAT, MinT, MaxT, T_range_) could be analysed as independent variables or not, we generated a correlation matrix using the R package *corrplot* (Wei & Simko 2021). Since almost all local climate variables were moderately to strongly correlated with each other, we elected to conduct a Principal Components Analysis (PCA) with all five variables. The PCA approach means that we cannot state explicitly that MAT or MAP or T_range_ are strong explanatory variables individually *per se*. Rather, the distinct biomes are defined clearly by climate variables in combination. The first two Principal Component axes (PC1 and PC2, respectively) explained 99.6% of the variation among the five climate variables; therefore, we used these two axes as local climate variables in analyses. The PC1 axis represents a continuum from cool and wet to extremely hot and dry. The PC2 axis represents a continuum from thermally stable with benign minimum temperatures to thermally variable with cold minimum temperatures.

Our first objective was to assess if and how the thermal tolerance thresholds and TTB of a broad range of plant species differed among the contrasting biomes, while our second objective was to test whether species from the more thermally variable alpine and desert biomes had wider thermal tolerance breadth than temperate biome species. To address both these objectives, we first calculated summary statistics (means and standard errors) for each thermal tolerance threshold metric by biome and species. Next, we fitted individual linear models for the four response variables and conducted analysis of variance (ANOVA). All response variables showed highly significant differences among biomes based on ANOVA; therefore, we then computed and reported pairwise differences using Tukey’s Honest Significant Difference method to compare pairs of biomes.

Our third objective was to assess whether variation in thermal tolerance thresholds and TTB could be explained by local climate variables, and whether these relationships were linear or not. To also account for variation in thermal tolerance thresholds and TTB due to phylogenetic relatedness, we tested the relationships between the thermal tolerance threshold variables and local climate variables using linear mixed effects regression models using the *Almer* function from the *evolvability* R package (Bolstad *et al*. 2014). The *Almer* function is built on the *lmer* function from the *lme4* R package (Bates *et al*. 2015) but incorporates a correlated random effects structure *A*, which we used to fit a phylogenetic variance–covariance matrix to account for phylogenetic relatedness among species. For each response variable (T_crit-hot_, T_crit-cold_, T_nucleation_, TTB), we fit four candidate models that included fixed effects of both climate variables, PC1 and PC2, as either linear or second-degree (quadratic) polynomial terms in combination (i.e. fixed effects in model 1 = PC1 + PC2; model 2 = PC1 + PC1^2^ + PC2; model 3 = PC1 + PC2 + PC2^2^; model 4 = PC1 + PC1^2^ + PC2 + PC2^2^). Each candidate model included three random effects: species (69 levels), growth form (10 levels), and a structured phylogenetic variance–covariance matrix. Variance explained by differences among species could therefore be separated from variance explained by phylogenetic relatedness. We calculated Akaike’s Information Criterion (AIC) values for each model fit by maximum likelihood to allow for model comparisons, and then calculated Akaike’s weights using the *MuMIn* R package (Bartoń 2023) to identify the most parsimonious (final) model for each thermal tolerance threshold response variable. We then refit the final model using restricted estimate maximum likelihood (REML) to provide unbiased parameter estimates. Variance proportions (as %) were reported for species, growth form, and phylogeny by squaring the random effects coefficients (standard deviations) and calculating the proportion of the random effect coefficient (e.g. species) out of the total (all random effects including the residual) variance. Goodness-of-fit was evaluated by calculating *R*^2^_GLMM_ values (Johnson 2014) to show variance explained by fixed effects (marginal *R*^2^; *R*^2^*m*) and the variance explained by both fixed and random effects (conditional *R*^2^; *R*^2^*c*). To test for spatial autocorrelation based on sampling location, we investigated the residuals of final models for each thermal tolerance threshold and fitted separate sets of models for each biome subset, following the model 1 structure. We then used Moran’s *I* test with inverse distance weights (1/d^2^) based on sampling latitude and longitude to test for significant spatial autocorrelation in all model residuals with the *spdep* package (Bivand 2022).

## Results

### Objective 1: Differences in thermal tolerance thresholds among the contrasting biomes

There were significant differences among biomes for all thermal tolerance thresholds (Fig. 2A-C). A full list of species and their individual thermal tolerance thresholds and breadth is presented in Table S1. Unexpectedly, desert species exhibited the most extreme cold thresholds T_crit-cold_ (–13.2 ± 0.3 °C), followed by alpine (–10.9 ± 0.4 °C) and then temperate species (–8.8 ± 0.3 °C), all of which differed significantly from one another (Fig. 2A; Tables S1 and S3). Similar to the results of T_crit-cold_, T_nucleation_ values were lowest in desert species (–16.1 ± 0.4 °C), followed by alpine (–10.8 ± 0.5 °C) and highest in temperate species (–7.5 ± 0.4 °C). Again, T_nucleation_ values among biomes differed significantly from one another (Fig. 2B; Tables S1 and S3). Interestingly, T_crit-cold_ values for alpine and temperate species were not significantly different from their T_nucleation_, whereas in desert species, ice formation occurred at significantly lower temperatures than T_crit-cold_ (Fig. 2A,B; Fig. S2A,B; Table S4). As predicted, desert species had significantly higher heat tolerance T_crit-hot_ (49.3 ± 0.5 °C), followed by temperate (46.5 ± 0.4 °C) and then alpine (43.1 ± 0.4 °C) species (Fig. 2C; Tables S1 and S3).

**Fig. 2.**
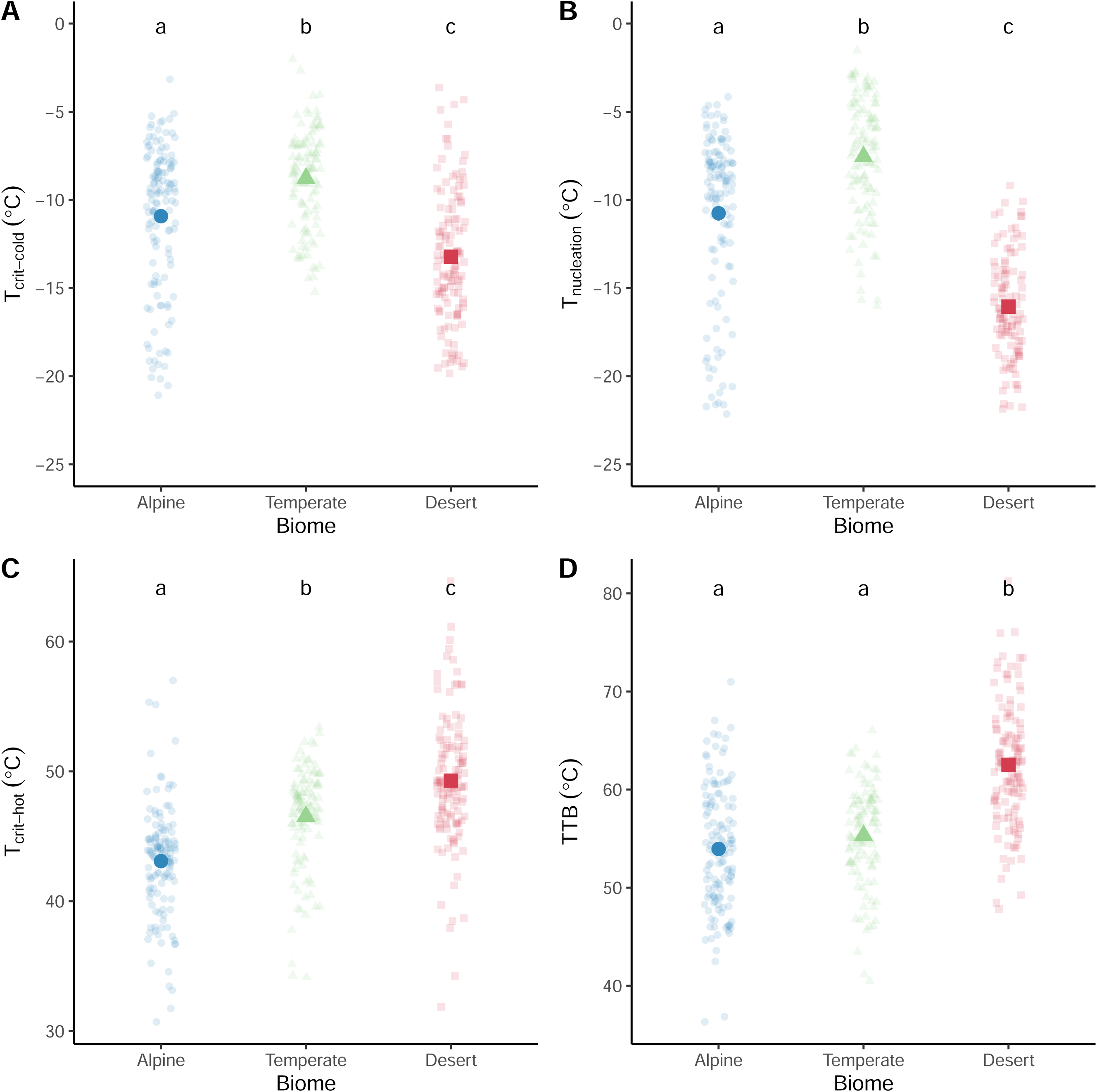
Thermal tolerance thresholds and breadth across the three biomes. (A) Cold tolerance threshold (T_crit-cold_), (B) ice nucleation temperature (T_nucleation_), (C) heat tolerance threshold (T_crit-hot_) and (D) thermal tolerance breadth (TTB = T_crit-hot_ - T_crit-cold_), where desert has *n* = 24 species, temperate has *n* = 23 species and alpine has *n* = 22 species. Small transparent points represent individual replicates, and large solid points represent biome means ± SE. Lowercase letters inset in each plot denote significant differences among biomes from post-hoc Tukey’s Honest Significant Differences tests within each thermal tolerance metric (see also Table S2).

### Objective 2: Test whether thermal tolerance breadth differs among biomes

Thermal tolerance breadth (TTB) was significantly wider in desert species (T_crit-cold_ - T_crit-hot_ = – 13.2 - 49.3 °C, TTB = 62.5 ± 0.6 °C), than temperate (T_crit-cold_ - T_crit-hot_ = –8.8 - 46.5 °C, TTB = 55.3 ± 0.5 °C) or alpine species (T_crit-cold_ - T_crit-hot_ = –10.9 - 43.1 °C, TTB = 53.9 ± 0.6 °C), but alpine and temperate species had similar TTB (Fig. 2D; Tables S1 and S3). There was considerable variation among species within biomes (Fig. S3). Within the desert biome, TTB was more similar across species (lower variability; TTB range = 56.2 - 70.3 °C; Δ = 14.1 °C), compared to the alpine biome, which had the greatest variation among species (TTB range = 45.9 - 63.1 °C; Δ = 17.2 °C) and the temperate biome with moderate variation (TTB range = 44.8 - 60.4 °C; Δ = 15.6 °C). The rank order of species’ TTB from widest to narrowest generally reflected their biome, where 17 of the top 20 widest TTBs were all desert species (Fig. S3). By contrast, 10 of the 20 narrowest TTBs were alpine species, and temperate species had generally moderate to narrow TTB (Fig. S3).

Consistent across all three biomes, TTB was much wider than the extreme temperature range calculated for the sampling sites; thus, most thermal tolerance thresholds extended beyond the minimum and maximum temperatures expected for each species in their local environment. The only notable exception was for the alpine biome species, where their cold threshold was closer to the mean minimum temperature of our alpine sites (Fig. 3).

**Fig. 3.**
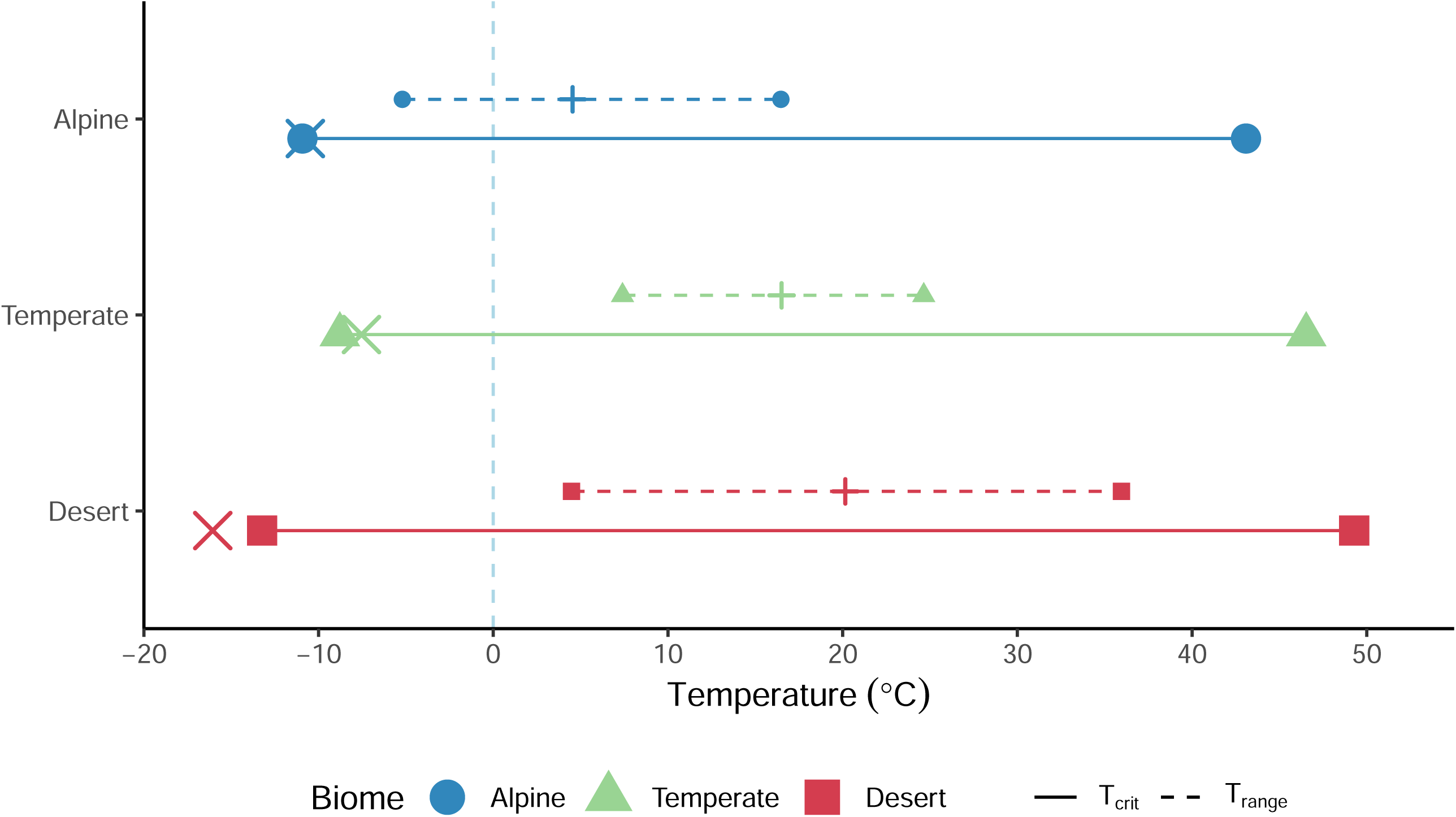
Mean thermal tolerance breadth (TTB = T_crit-hot_ -T_crit-cold_) for the alpine, temperate and desert biomes, relative to the mean thermal range of the local environments (T_range_ = MaxT - MinT). Red squares and lines represent desert, green triangles and lines represent temperate and blue circles and lines represent alpine biomes. Cross (×) symbols represent the mean T_nucleation_ and plus (+) symbols represent the mean annual temperature for each biome. Solid lines connect the mean T_crit-cold_ to the mean T_crit-hot_ (large symbols) for each biome (length of line = TTB). Dashed lines connect the mean MinT to the mean MaxT (small symbols) for each biome (length of line = T_range_). The light blue vertical dashed line represents 0 °C. Each species has *n* = 5-6 replicates for each thermal tolerance threshold value.

### Objective 3: Local climate drivers of variation in thermal tolerance thresholds and breadth

Most of the local climate variables were strongly correlated with each other (Fig. 4A); therefore, by conducting a PCA, 99.6% of the climate space among the three biomes could be described along two axes (PC1 and PC2; Table S5; Fig. 4B). PC1 explained 77.2% of the variation among the five climate variables and was strongly loaded negatively by MAP and positively by MAT and MaxT (Fig. 4B,C). Negative PC1 values can be reasonably interpreted as cooler and wetter climate, while positive PC1 values indicate a hotter (mean and extreme temperatures) and drier climate (Fig. 4B). PC2 explained 22.4% of the variation among the five climate variables and was strongly loaded positively by T_range_ and negatively by MinT (Fig. 4B,D). Positive PC2 values describe a climate with colder extremes and greater range in thermal extremes across the year, while negative PC2 values indicate a climate with less extreme cold temperatures and greater thermal stability through the year (Fig. 4B). No evidence for spatial autocorrelation was detected from residuals in any model (Table S6).

**Fig. 4.**
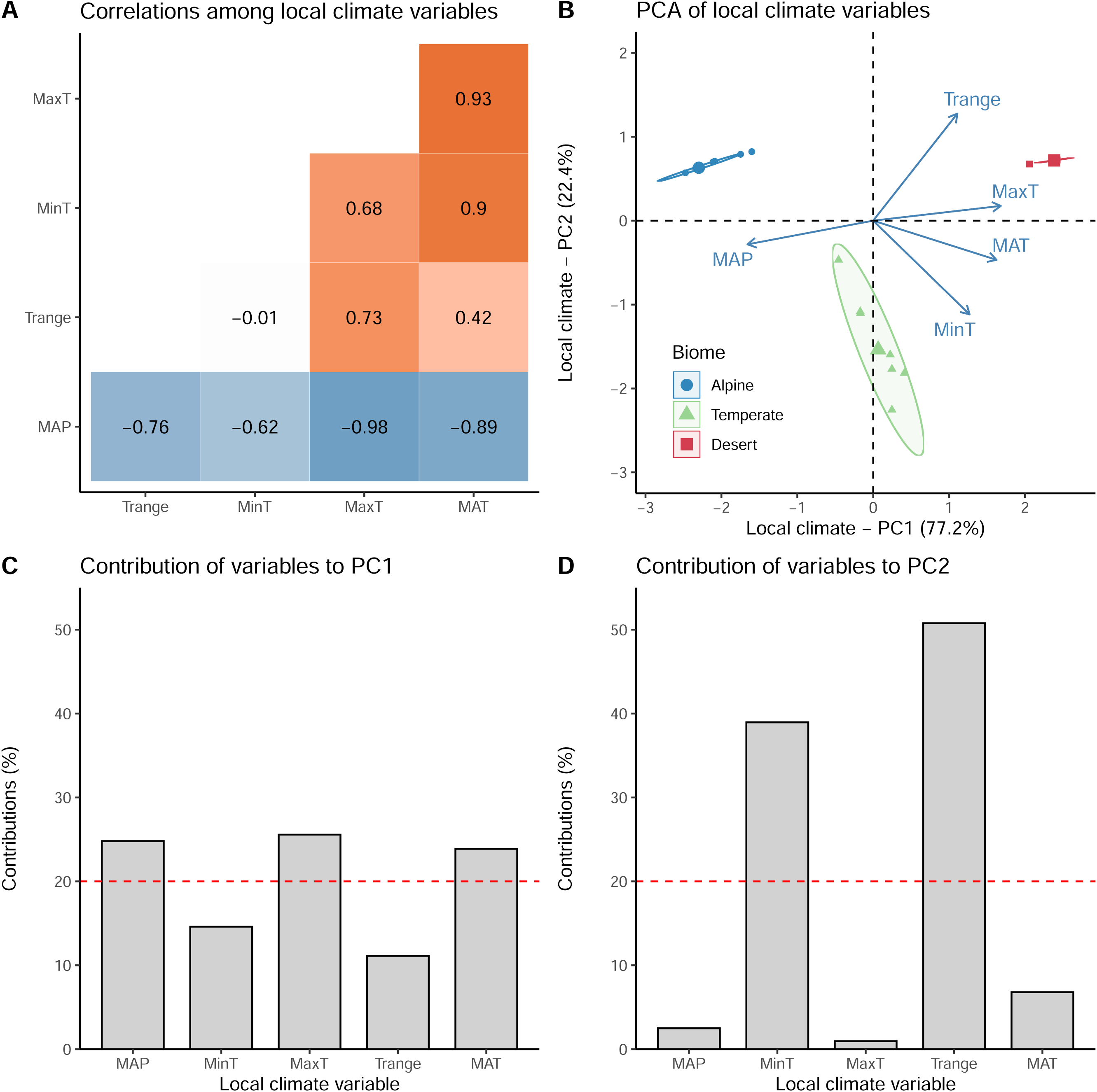
Correlations and Principal Component (PC) composition of local climate variables (MAP = mean annual precipitation, MAT = mean annual temperature, MinT = mean minimum temperature of the coldest month, MaxT = mean maximum temperature of the warmest month, T_range_ = annual thermal range (MaxT - MinT)). (A) Correlation matrix of local climate variables, (B) Principal Components Analysis biplot of PC1 and PC2 showing biome position in climate PC space, and contribution of individual local climate variables to (C) PC1 and (D) PC2.

We investigated the relationships between the Principal Components of climate variables at the local site of plant sampling and each of the thermal tolerance metrics. Model comparisons (AIC and weights) for model selection are shown in Table S7. We found that T_crit-cold_ had a significant non-linear relationship with PC1 but no significant relationship with PC2 (Table 1, Fig. 5A,B). That is, the highest cold tolerance (i.e. most negative) was detected in plants in the desert, followed by those in alpine, then those in temperate biomes. T_nucleation_ had a significant (negative) linear relationship with PC1 and significant non-linear relationship with PC2 (Table 1, Fig. 5C,D). That is, the greatest freezing tolerance (i.e. ice nucleation at colder temperatures) was detected in plants growing in hot, dry climate (desert). Plants from more thermally stable and less extreme temperate climate were less tolerant of freezing, but freezing tolerance increased sharply in climates with colder extremes and greater range in thermal extremes across the year (both desert and alpine) (Fig. 5D). Species from environments with greater seasonality (i.e. climate variability) therefore froze at lower temperatures than species in environments where temperature is more stable (Fig. 5D).

**Fig. 5.**
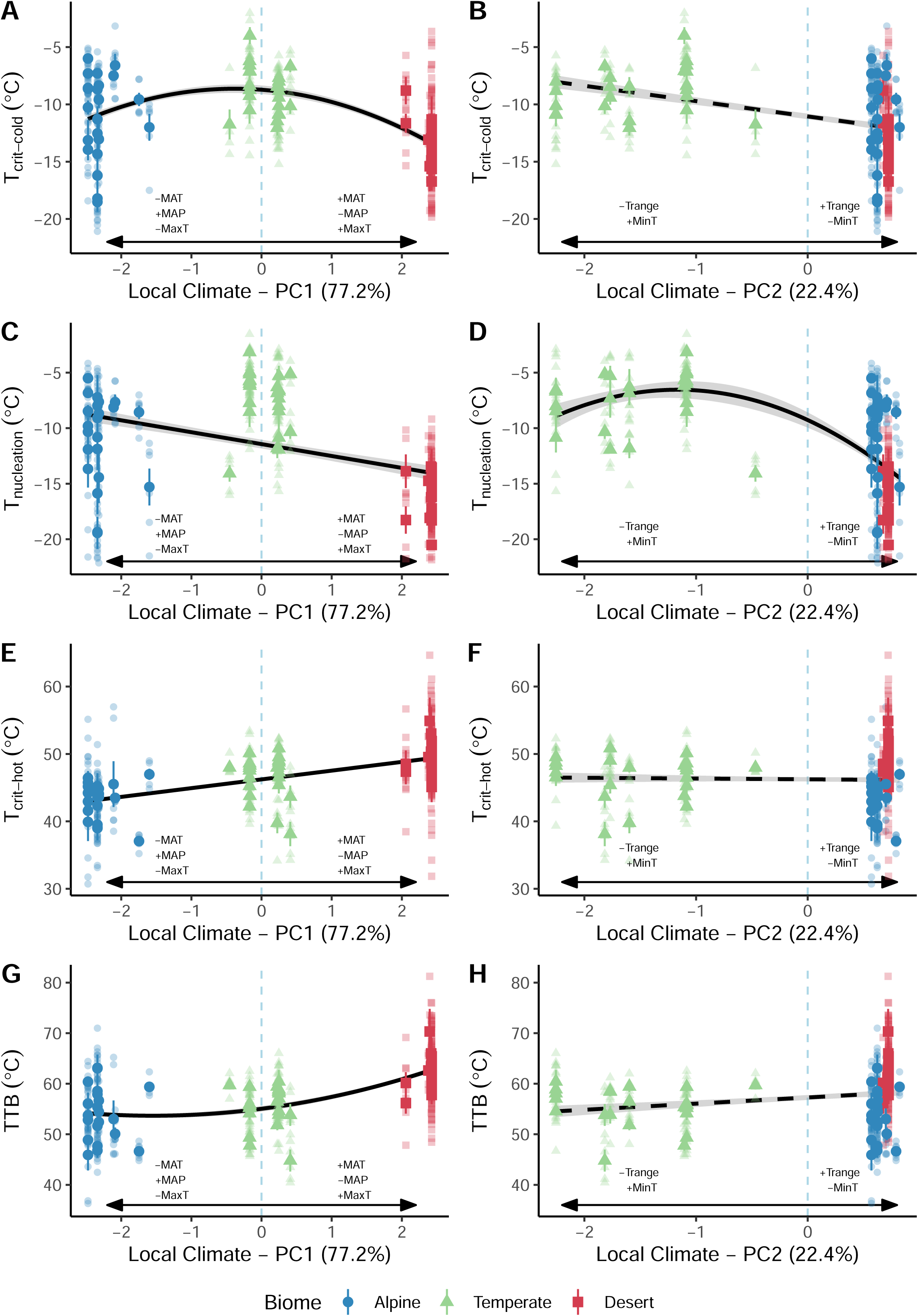
Local climate variables (Principal Components, PC1 and PC2; see Fig. 4) that predict variation in thermal tolerance thresholds across all biomes. Regressions between (A) T_crit-cold_ and PC1, (B) T_crit-cold_ and PC2, (C) T_nucleation_ and PC1, (D) T_nucleation_ and PC2, (E) T_crit-hot_ and PC1, (F) T_crit-hot_ and PC2, (G) TTB and PC1 and (H) TTB and PC2. Arrows and annotations indicate climate variable loadings on PC1 and PC2 axes: positive PC1 values = hotter and drier climate (+MAT, –MAP, +MaxT); negative PC1 values = cooler and wetter climate (–MAT, +MAP, - MaxT). Positive PC2 values = greater range in thermal extremes across the year and colder extremes (+T_range_, –MinT) negative PC2 values = greater annual thermal stability and less extreme cold temperatures (–T_range_, +MinT), see Fig. 4 and Table S5. Large solid points are species-level means ± SE and small transparent points are individual replicates. Points are coloured by biome and regressions are predicted model fits ± SE from the best models reported in Table 1, where solid regression lines represent significant relationships.

**Table 1.** Results of linear mixed-effects regression models that tested the relationships between local climate factors (summarised as Principal Components, PC1 and PC2) and thermal tolerance thresholds.

| Cold tolerance threshold: $T_{crit-cold}$ (°C) | | | | | | | |
| --- | --- | --- | --- | --- | --- | --- | --- |
| Fixed | Coefficient ± SE | *t*-value | *p*-value | Random | Coefficient | $R^2$ type * | $R^2$ value |
| Intercept | $-10.488 \pm 1.167$ | -8.985 | <0.001 | Growth form | 1.308 | $R^2_m$ | 0.195 |
| PC1 | $-15.179 \pm 6.606$ | -2.299 | 0.022 | Species | 1.829 | $R^2_c$ | 0.515 |
| PC1<sup>2</sup> | $-38.459 \pm 15.449$ | -2.487 | 0.013 | Phylogeny | 0.081 | $R^2_c - R^2_m$ | 0.320 |
| PC2 | $0.451 \pm 0.774$ | 0.582 | 0.561 | Residual | 2.768 | | |
| Freezing tolerance: $T_{nucleation}$ (°C) | | | | | | | |
| Fixed | Coefficient ± SE | *t*-value | *p*-value | Random | Coefficient | $R^2$ type * | $R^2$ value |
| Intercept | $-11.432 \pm 0.585$ | -19.551 | <0.001 | Growth form | 1.149 | $R^2_m$ | 0.430 |
| PC1 | $-0.935 \pm 0.205$ | -4.567 | <0.001 | Species | 2.516 | $R^2_c$ | 0.725 |
| PC2 | $-49.355 \pm 7.135$ | -6.917 | <0.001 | Phylogeny | 0.000 | $R^2_c - R^2_m$ | 0.295 |
| PC2<sup>2</sup> | $-15.707 \pm 6.469$ | -2.428 | 0.016 | Residual | 2.670 | | |
| Heat tolerance threshold: $T_{crit-hot}$ (°C) | | | | | | | |
| Fixed | Coefficient ± SE | *t*-value | *p*-value | Random | Coefficient | $R^2$ type * | $R^2$ value |
| Intercept | $46.360 \pm 0.415$ | 111.662 | <0.001 | Growth form | 0.518 | $R^2_m$ | 0.250 |
| PC1 | $1.318 \pm 0.184$ | 7.148 | <0.001 | Species | 2.236 | $R^2_c$ | 0.446 |
| PC2 | $-0.166 \pm 0.328$ | -0.504 | 0.615 | Phylogeny | 0.000 | $R^2_c - R^2_m$ | 0.196 |
|  |  |  |  | Residual | 3.856 |  |  |
| Thermal tolerance breadth: TTB (°C) |  |  |  |  |  |  |  |
| Fixed | Coefficient ± SE | *t*-value | *p*-value | Random | Coefficient | $R^2$ type * | $R^2$ value |
| Intercept | $57.440 \pm 0.928$ | 61.920 | <0.001 | Growth form | 2.035 | $R^2_m$ | 0.273 |
| PC1 | $65.441 \pm 10.543$ | 6.207 | <0.001 | Species | 3.146 | $R^2_c$ | 0.550 |
| PC1<sup>2</sup> | $49.280 \pm 24.352$ | 2.024 | 0.044 | Phylogeny | 0.000 | $R^2_c - R^2_m$ | 0.276 |
| PC2 | $-1.102 \pm 1.216$ | -0.906 | 0.365 | Residual | 4.789 | | |

Note: PC1 represents an axis from cooler and wetter climate (negative values) to hotter mean and extreme temperatures and drier climate (positive values). PC2 represents an axis from a climate with less extreme cold temperatures and greater thermal stability through the year (negative values) to a climate with colder extremes and greater range in thermal extremes across the year (positive values). \* $R^2_m$ represents the variance explained by all fixed effects, $R^2_c$ represents the variance explained by all fixed and random effects, and $R^2_c - R^2_m$ represents the variance explained by all random effects.

Heat tolerance had a significant linear relationship with PC1 but not PC2 (Table 1). T_crit-hot_ was highest in plants from hot, dry climate and it declined as the climate became increasingly cool and wet (Fig. 5E,F). Thermal tolerance breadth had a significant non-linear relationship with PC1 but was not associated with PC2 (Table 1, Fig. 5G,H). TTB was highest in hot, dry climates and it declined non-linearly as climates became cooler and wetter, where TTB was not different between alpine and temperate species (Fig. 5G). For all thermal tolerance thresholds, among-species differences explained a substantial proportion of variance (variance proportions due to species from Table 1 as percentage: T_crit-cold_ = 26.3%, T_nucleation_ = 42.8%, T_crit-hot_ = 24.8%, TTB = 26.8%). Plant growth form explained some variance, which was greatest for TTB (variance proportions due to growth form from Table 1 as percentage: T_crit-cold_ = 13.4%, T_nucleation_ = 8.9%, T_crit-hot_ = 1.3%, TTB = 11.2%). Among-species differences were not attributable to phylogenetic relatedness (Table 1). All four phylogenetic signal metrics consistently identified no evidence of phylogenetic signal in the thermal tolerance thresholds or breadth (Table S8).

## Discussion

The climate variability hypothesis (CVH) predicts that species will evolve wider thermal tolerance breadth (TTB) in environments with more variable temperatures and be selected for thermal specialisation (narrow TTB) in thermally stable environments. The CVH has been tested in a few plant systems across different contexts, with limited support (Molina-Montenegro & Naya 2012; Cuesta *et al*. 2020; Wooliver, Tittes & Sheth 2020; Querns *et al*. 2022; Chiono & Paul 2023). For example, two studies on *Mimulus guttatus* did not find support for stronger seasonality being associated with TTB based on thermal performance curves of relative growth rate (Querns *et al*. 2022; Chiono & Paul 2023). By contrast, another study across the geographic range of *Mimulus cardinalis* found that TTB of relative growth rate was indeed broader in populations found in more seasonal climates (Wooliver, Tittes & Sheth 2020). Here we explored the CVH in diverse plants by applying high-throughput measurements of heat and cold tolerance thresholds of photosystem II concurrently to determine TTB of species growing in two opposite extremes and thermally variable biomes (desert and alpine) and one benign, thermally stable biome (temperate). Overall, we found that mean annual and mean maximum temperatures, and mean annual precipitation had a stronger relationship with TTB than did thermal variability and mean minimum temperatures, which provides only partial support for the CVH. Below we explore our findings in terms of physiology and the ecological implications of these results for species persistence in a changing climate.

### Heat and cold tolerance thresholds vary significantly among biomes (Objective 1)

Based on the CVH framework, we predicted that the thermal tolerance thresholds of alpine and desert species would be skewed to their respective biome extremes; that is, colder threshold temperatures in the alpine and hotter threshold temperatures in the desert, with temperate species having both milder cold and heat tolerance thresholds. Here we found partial support for this prediction. Temperate species had heat tolerance intermediate between alpine and desert but were less cold tolerant than either alpine or desert; desert species were more heat tolerant than alpine and temperate species but were also more tolerant of cold and freezing than alpine species.

The apparently counterintuitive result that desert species were more tolerant of cold and resistant to freezing (more extreme T_crit-cold_ and T_nucleation_ values; Fig S2) than either alpine or temperate species may be partially explained by the physiological response to drought conditions. That is, plants exposed to drought and/or freezing share a similar physiological response to cell dehydration (Anisko & Lindstrom 1996; Blake & Hill 1996; Lintunen, Hölttä & Kulmala 2013). Plants can use osmotic adjustments to offset water loss from cells along the water potential gradient caused by both drying and freezing (Siminovitch & Cloutier 1983; Larcher 2003). In response to freezing conditions, plants typically either tolerate ice formation by coping with cellular dehydration (Scholz *et al*. 2012) or avoid ice formation by supercooling, which refers to preventing ice formation below the equilibrium freezing temperature of the leaf tissue (Pearce 2001). Here, the desert species that we studied generally had ice nucleation temperatures (T_nucleation_) that were much lower than their T_crit-cold_, from which we can infer a high supercooling capacity that delays freezing (Fig. S2). There are several examples that demonstrate the link between freezing and dehydration. Extreme T_nucleation_ found in Andean alpine species can be attributed to low soil moisture rather than low temperatures (Sierra-Almeida, Reyes-Bahamonde & Cavieres 2016). Drought also improves both heat and cold tolerance in an alpine grass (Sumner *et al*. 2022), and drought can increase heat tolerance by raising intracellular sugar content, which stabilises thylakoid membranes (Hüve *et al*. 2006; Brestic *et al*. 2012). Therefore, we postulate that the extreme tolerance in these desert species reflects a secondary effect of their adaptations to aridity.

Alpine plants generally tolerated cold and freezing very well, as expected. Retaining high cold tolerance remains necessary for protecting investment in valuable photosynthetic tissue due to exposure to subzero temperatures without insulative snow during growing seasons (Slatyer, Umbers & Arnold 2022). The relatively high heat tolerance of some of our alpine species may seem surprising; however, these heat tolerance ranges are similar to recent measures of heat tolerance in other alpine species (43-55 °C, Sumner & Venn 2022; Arnold *et al*. 2024; Harris *et al*. 2024; Morales *et al*. 2025). Alpine plants can be exposed to very high temperatures, especially those with prostrate growth forms (Körner 2003). Leaf temperature of European alpine species can reach 50 °C in summer on calm sunny days, and leaf heat tolerance thresholds in these plants can exceed 50 °C (Buchner & Neuner 2003). Maximum leaf temperature has been measured at nearly 40 °C in early summer in Australian alpine species (Bird *et al*., unpublished data) and may yet be much higher in hot and still conditions in late summer. Therefore, despite the apparently high heat tolerance thresholds, these could be breached in alpine species if heat interacts with other environmental stress, such as high light, which can cause heat tolerance to decrease in some European alpine species (Buchner & Neuner 2003; Buchner *et al*. 2015).

The tolerance of temperate plants was generally less than that of plants from the more extreme biomes, reflecting the more stable and less extreme climate of the biome. These temperate species tolerated temperatures well outside their typical range. Nevertheless, under warmer conditions in the future, vapour pressure deficit, which drives higher evaporative demand, will increase in wet temperate and tropical rainforests (Binks *et al*. 2023). Plants growing in these conditions will have greater water loss, and if water becomes limiting, transpirational cooling capacity will diminish, consequently pushing leaf temperature closer to heat tolerance thresholds (Zhou *et al*. 2023).

### Thermal tolerance breadth is wider in more thermally variable alpine and desert biomes and exceeds the thermal range of each biome (Objective 2)

The thermal tolerance breadth (TTB) values based on PSII tolerance that we report here are notably wide––wider than the thermal ranges in each environment and wider than many other measurements of TTB in other organisms. Across a range of marine and terrestrial ectotherm taxa, the widest recorded TTB was 60 °C in terrestrial arthropods (Sunday, Bates & Dulvy 2011), while the widest TTB in our dataset was 70.3 °C in the desert species *Rhagodia spinescens*, with several other desert species also exceeding 60 °C. Within each biome, we found species with a range of wide and narrow TTB (Fig. S3). For example, several alpine species across growth forms had comparatively narrow TTB, such as the snow bed species *Psychrophila introloba* (TTB = 48.9 °C) and the tree line species *Eucalyptus pauciflora* subsp. *niphophila* (TTB = 47.7 °C). The narrow TTB of some alpine species was mainly driven by their relatively low heat tolerance thresholds (T_crit-hot_ ≍ 40 °C), whereas other co-occurring species, *Hovea montana* and *Epacris paludosa* had TTB values similar to those of many desert species (∼63 °C), which was driven by their extreme cold tolerance (T_crit-cold_ ≍ –18 °C). The notable within-biome variation in TTB might be due to variation in microclimatic niche among species, which can override broad-scale climate drivers of physiological tolerance (Curtis *et al*. 2016; Curtis, Knight & Leigh 2019; Aparecido *et al*. 2020).

Although thermal seasonality was greater in the alpine than in the temperate regions sampled here, the average TTB for these two biomes was remarkably similar (Fig. 2D). Temperate species appeared to tolerate colder temperatures than expected, and it is possible that their relatively wide thermal tolerance is a legacy of exposure to climate extremes through geological time (Byrne *et al*. 2008). The climate legacy effect on thermal tolerance is suggested to be stronger for cold than for heat tolerances across both animal and plant taxa (Bennett *et al*. 2021), which might be an additional explanation as to why some temperate and desert species have greater freezing tolerance than expected based on their current climate envelopes. Although climate metrics based on air temperatures for our alpine biome confirm that thermal range and seasonality are large in this biome, these mountain plants can spend ∼4 months under the insulating cover of snow, which stabilises plant temperature (Briceño *et al*. 2014). The dampening of climate extremes at a local scale (i.e. microclimate) might also contribute to similar TTB values between alpine and temperate species.

We find that when considering climate variability from a discrete biome perspective, our results provide only partial support for the CVH. That is, plants from the climatically variable desert biome have the widest TTB, but temperate and alpine plants do not have different TTB despite the greater daily and seasonal climate variability in the alpine environment. Investigating the climatic conditions occurring in each biome provides further insight into the drivers behind each thermal tolerance threshold and TTB.

### Thermal tolerance thresholds and breadth are driven by different climatic variables (Objective 3)

Thermal tolerance is difficult to predict based on individual bioclimatic parameters, such as mean annual temperature and mean annual precipitation (Perez & Feeley 2021). Here, the Principal Component axis that combined MAT, MAP and MaxT explained considerable variation in T_crit-cold_ and T_nucleation_. As discussed above, the much lower T_nucleation_ than T_crit-cold_ in desert species suggests that T_nucleation_ is more strongly associated with aridity than is T_crit-cold_, supporting the idea of shared mechanisms of tolerance to freezing and dehydration under drought. Although plants growing in extremely hot and dry environments do have higher heat tolerance, the high correlation of MAP with both MAT and MaxT means that the role of temperature versus precipitation cannot be completely separated. For a range of common-grown Australian *Acacia* species from widespread climatic and geographic origins, precipitation did not strongly drive variation in T_crit-hot_ (Andrew *et al*. 2023). Even so, water availability and temperature are intrinsically linked in terms of plant stress responses. Water limitation can drastically reduce stomatal conductance and therefore cooling capacity during exposure to heat, increasing leaf temperature and affecting heat tolerance thresholds (Cook *et al*. 2021).

Local climate, particularly MAT and MaxT, seems to play a significant role in driving variation in T_crit-hot_ in our field study, which contrasts with a common garden study of diverse species, for which origin climate generally had weak predictive power for heat tolerance (Perez & Feeley 2021). It is important to reiterate that MinT and MaxT as analysed here are mean values. Daily or hourly extreme temperatures can far exceed MinT and MaxT for short periods; therefore, the apparently wide buffer that TTB may provide could occasionally be much narrower than these patterns suggest. However, it is also important to recognise that many plants have the capacity to regulate their leaf temperature. A controlled environment study on a subset of species studied here found that alpine and desert plants cooled their leaves by 1-3 °C below air temperature during a 38 °C heat event, while temperate plants were frequently 1-2 °C above air temperature (Arnold *et al*. 2025). Strong thermal decoupling has been found in less heat-tolerant alpine plant species as an effective heat avoidance strategy (Morales *et al*. 2025). When water is not limiting, some species can thermoregulate to prevent leaves from reaching damaging temperatures, but that strategy may be nullified during hotter droughts (Cook *et al*. 2021; Arnold *et al*. 2025). Underpinning thermoregulation strategies are combinations of leaf structural traits (leaf density, thickness, water content) and stomatal behaviour that also vary among species and across biomes (Arnold *et al*. 2025).

A large proportion of the variation in thermal tolerance thresholds in our study was also attributable to among-species differences that were unrelated to phylogeny. The contribution of species differences could reflect their different leaf traits, particularly those contributing to variation in leaf temperature, the signal of which may be amplified in the field (Leigh *et al*. 2017). Recent studies have shown that a major driver of variation in heat tolerance is leaf temperature, rather than air temperature (Perez & Feeley 2020; Cook *et al*. 2021). Leaves can avoid heat stress through different mechanisms that decouple leaf from air temperature, such as anatomical traits (Leigh *et al*. 2012; Leigh *et al*. 2017; Tserej & Feeley 2021; Arnold *et al*. 2025), leaf inclination (Ball, Cowan & Farquhar 1988) and transpirational cooling (Drake *et al*. 2018; Deva *et al*. 2020), all of which can vary among co-existing species (Urban *et al*. 2017; Marchin *et al*. 2022).

The capacity for different species to respond to increasingly extreme and variable temperatures by changing their TTB is only recently emerging. We found that TTB increased non-linearly (hyper-allometrically) with higher MAT and MaxT and lower MAP (Fig. S4). That is, plants exposed to the high mean temperatures, coupled with aridity and heat extremes in deserts had disproportionally wider TTB than temperate or alpine plants. TTB of high elevation species from temperate and tropical regions appears to differ only at the end of the temperate growing season, due to an increase in cold acclimation (Sklenář *et al*. 2023). In a commongarden experiment with seedlings from the same biomes that were tested here, TTB narrowed when exposed to a combination of hot days and freezing nights in controlled climate chambers (Harris *et al*. 2024). However, that study found a minimal effect of biome, which contrasts with the significant biome-based differences in thermal tolerance thresholds observed here for plants *in situ*. The distinct responses between these contexts imply that life stage and current growing conditions could be important modifiers of thermal tolerance.

The effects of immediate growth conditions can outweigh other environmental and genetic sources of variation in trait changes in plants grown in common gardens (Arnold *et al*. 2024). In alpine species at least, both cold and heat tolerance of PSII can vary widely among genetically similar populations distributed along an elevational gradient, suggesting intraspecific variance in thermal tolerance is driven by acclimation to local conditions at a fine scale (Danzey *et al*. 2024). Whether PSII thermal tolerance is heritable and under selection in wild plants remains largely unknown, but evidence so far suggests it is not (Arnold *et al*. 2024). PSII thermal tolerance can be highly heritable in rice species (Robson *et al*. 2023). Yet, evidence from wheat shows T_crit-hot_ in field-planted crops is strongly dependent on growth temperature during the days preceding anthesis, and that T_crit-hot_ is far less heritable in the field than in controlled environments (Coast *et al*. 2022). Together, these studies and our findings here suggest that PSII thermal tolerance thresholds and TTB will be dependent both on the physical characteristics that differentiate species and local climate factors at fine spatial and temporal scales.

## Conclusions

While there were substantial differences in thermal tolerance thresholds among biomes, we found limited support for the climate variability hypothesis. Rather than climate variability, warmer mean annual temperature, maximum temperature, and lower precipitation were stronger drivers of wider TTB, particularly in desert plants, which were more tolerant of both extreme heat and cold than alpine or temperate plants. Thermal tolerance breadth of PSII in leaves was extremely wide in all biomes––wider than the typical air temperature extremes to which plants are naturally exposed in these biomes. Sustained extreme temperatures within the bounds of thermal tolerance breadth could still have far-reaching effects, depending on species life history, morphological traits and plasticity, as well as microclimatic and temporal heterogeneity. While local climate partially explains variability in plant thermal tolerance thresholds and breadth, incorporating additional traits, leaf temperature and microhabitat information through time are important next steps to better understand the vulnerability of different vegetation communities to climate change and extreme climatic events.

## Acknowledgements

We acknowledge the Traditional Custodians of Country throughout Australia, especially across the lands on which this study took place, and recognise their continuing connection to these lands, waters and natural communities. We pay our respects to Aboriginal and Torres Strait Islander cultures and to Elders past, present and emerging. We thank the National Parks and Wildlife Service for assistance and support, with special thanks to M. Hams, S. Doak and R. Bjork at Gundabooka National Park, M. Schroder at Kosciuszko National Park, and J. Erskine, J. Lemmon and P. Nagle at Royal National Park and Illawarra State Conservation Area. We thank the following people for their assistance in the field: C. Bellotto, Z. Brown, A. Crimmins, C. Cunningham, P. Cunningham, L. Danzey, M. Davies, L. Fuller, J.M. Hoskins, T. Salisbury, E. Rogers and K. Spooner. This study was conducted under approval from the NSW Department of Planning, Industry and Environment (Scientific Licence SL102330). We thank the anonymous reviewers and editors that provided constructive feedback that improved this manuscript. The work was supported by an Australian Research Council Linkage Project grant (LP180100942) in partnership with the Save our Species program, New South Wales Government Department of Planning, Industry and Environment.

## Conflict of interest statement

The authors have no conflicts of interest to disclose.

## Author Contributions

Verónica F. Briceño, Alicia M. Cook, Stephanie K. Courtney Jones, Adrienne B. Nicotra, León A. Bravo and Andy Leigh designed the study. Pieter A. Arnold, Rachael V. Gallagher, Verónica F. Briceño and Alicia M. Cook collated and analysed the data. Verónica F. Briceño and Pieter A. Arnold led the writing and revisions of the manuscript. All authors contributed to intellectual conceptualisation, data collection, interpretation of results and writing. All authors meet criteria for authorship.

## Data availability statement

Data and code supporting this study are available in the figshare repository at https://doi.org/10.6084/m9.figshare.29964764 (Briceño *et al*. 2025).

## Notes

https://doi.org/10.6084/m9.figshare.29964764

### Competing Interest Statement

The authors have declared no competing interest.

### Summary of Updates

Updated to match final text revisions at Journal of Ecology. Title shortened and some typos corrected.

