## Supplementary material for "Drivers of thermal tolerance breadth of plants across contrasting biomes": Fig. S1

**Supporting Information**

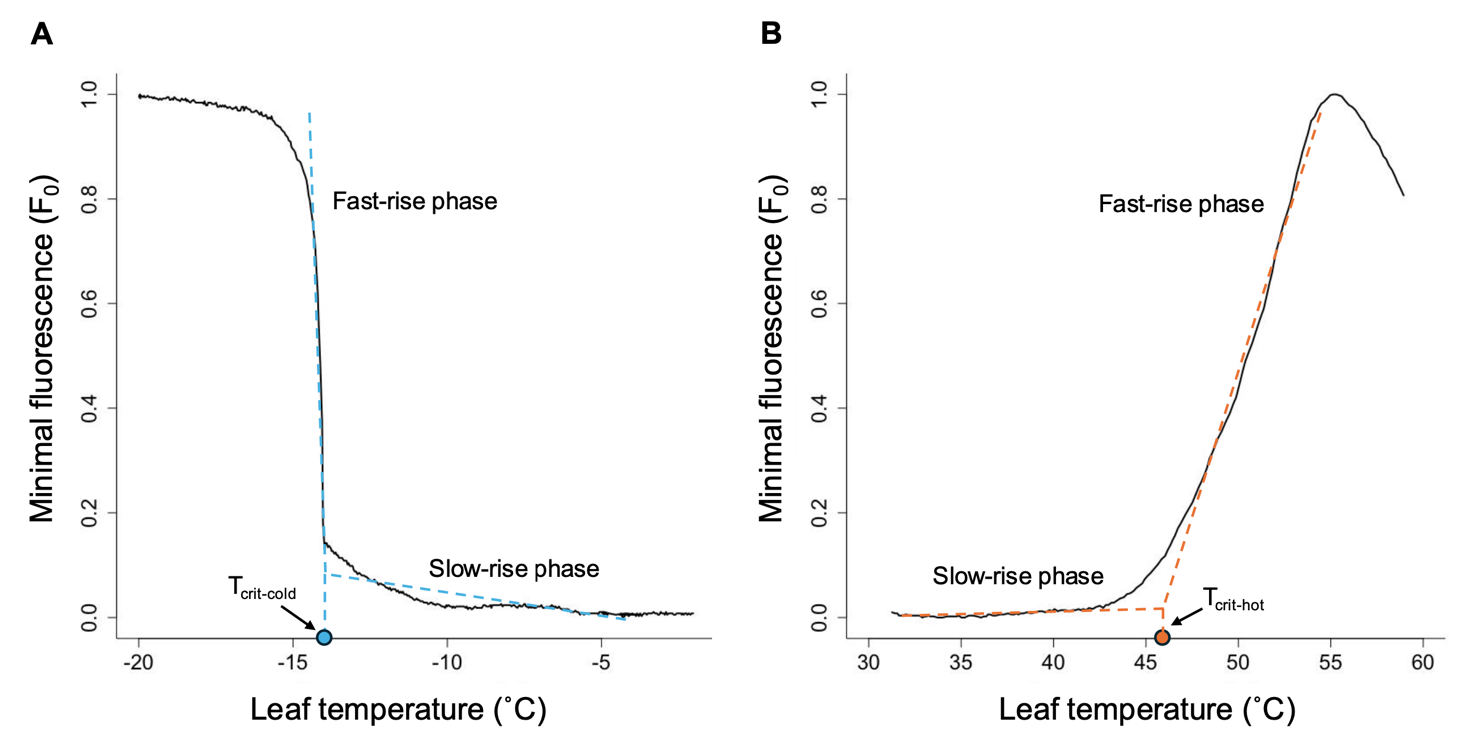

**Figure S1.** Example T-F_0_ curves for deriving T_crit-cold_ and T_crit-hot_ values. T-F_0_ refers to the temperature-dependent (T) change in minimal chlorophyll fluorescence (F_0_), when temperature is either increased or decreased at a steady rate. (A) T_crit-cold_ and (B) T_crit-hot_ were each determined as the point of transition between slow-rise and fast-rise phases of F_0_ that occurs during steady cooling or heating. Coloured dashed lines illustrate breakpoint regressions with vertical extension to the *x*-axis to show the leaf temperature that represents T_crit-cold_ (blue) and T_crit-hot_ (orange).

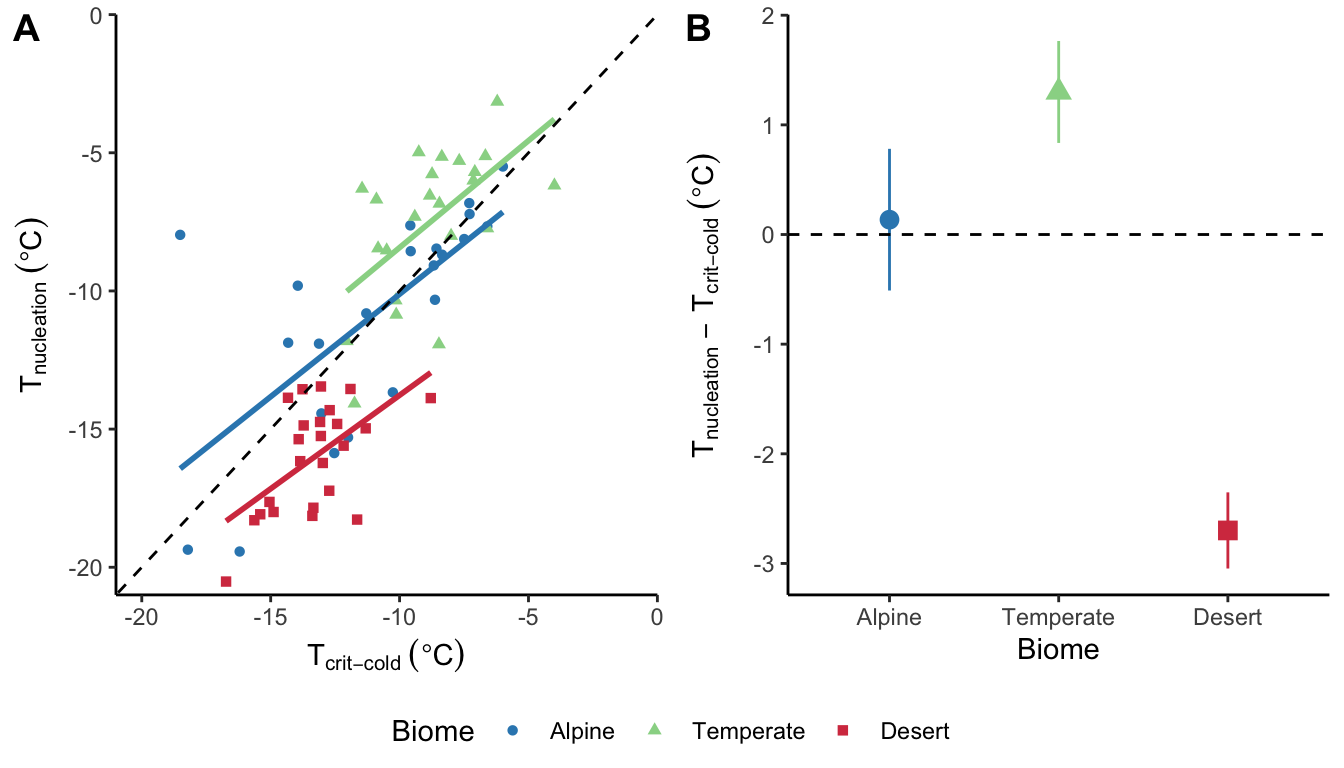

**Fig. S2.** (A) Relationship between the lower critical threshold T_crit-cold_ (°C) and ice nucleation (freezing point) temperature T_nucleation_ (°C). (B) Mean differences between T_nucleation_ and T_crit-cold_ (°C) across the three biomes, where negative values indicate a lower freezing point than critical threshold. The black dashed line is an isometric (1:1) line. Data points are species-level means. Linear regressions are fitted to each biome separately, which illustrates that species from the desert have a significantly lower T_nucleation_ than T_crit-cold_ (for model output see Table S3).

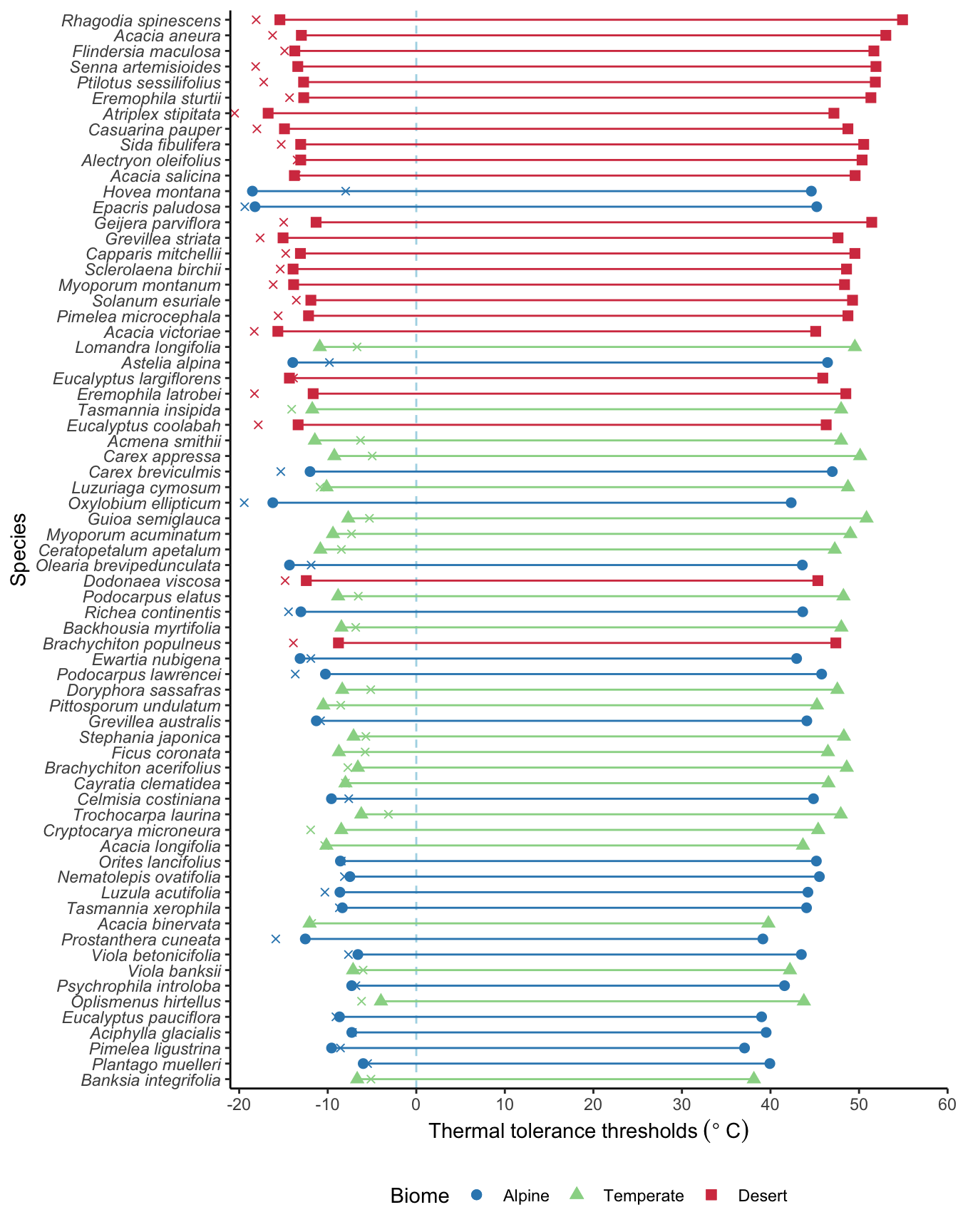

**Fig. S3.** Thermal tolerance breadth (TTB = T_crit-hot_ – T_crit-cold_) for 69 species across three biomes. Species are sorted top to bottom from widest to narrowest TTB, with lines connecting the mean T_crit-cold_ to the mean T_crit-hot_ for each species (length of line = TTB). Red squares and lines represent desert species (*n* = 24 species), green triangles and lines represent temperate species (*n* = 23 species), and blue circles and lines represent alpine species (*n* = 22 species). Cross symbols (×) represent the mean T_nucleation_ for each species. The light blue vertical dashed line represents 0 °C. Each species has *n* = 5-6 replicates for each thermal tolerance threshold value. The rank order of species’ TTB from widest to narrowest (top to bottom) generally reflects their biome, where 17 of the top 20 widest TTBs were all desert species. By contrast, 10 of the 20 narrowest TTBs were alpine species, and temperate species had generally moderate to narrow TTB.

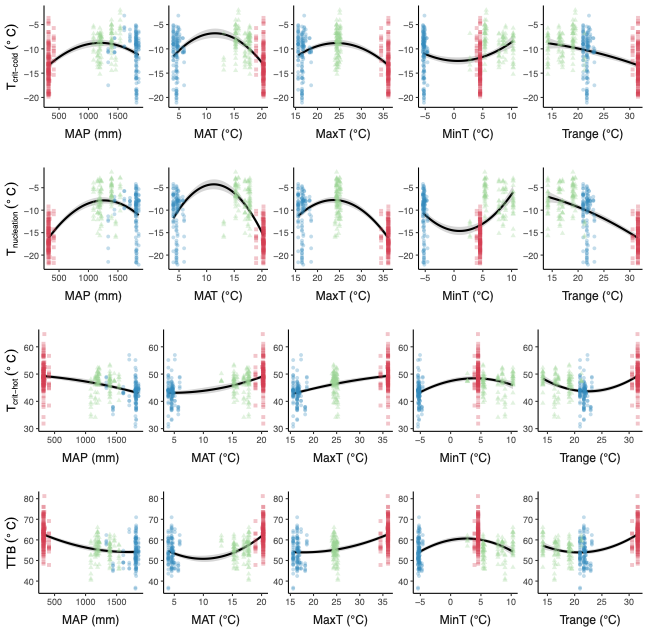

**Fig. S4.** Thermal tolerance thresholds for 69 species across three biomes plotted against individual local climate variables. Points are individual replicates coloured by biome (red squares = desert, green triangles = coastal temperate, blue circles = alpine) and regressions are quadratic fits for visualisation purposes only (analyses use PCA rather than individual climate variables due to strong correlations). MAP = Mean Annual Precipitation (mm), MAT = Mean Annual Temperature (°C), MaxT = Mean Maximum Temperature (°C), MinT = Mean Minimum Temperature (°C), and Trange = Annual Thermal Range (MaxT – MinT; °C).

**Table S1.** Selected species in each biome, family, growth form and mean thermal tolerance thresholds.

| Species | Family | Growth form | T_crit-cold_ (°C) | T_nucleation_ (°C) | T_crit-hot_ (°C) | TTB  (°C) |
| --- | --- | --- | --- | --- | --- | --- |
| Alpine |  |  |  |  |  |  |
| *Aciphylla glacialis* | Apiaceae | Herb | –7.3±0.4 | –7.2±0.3 | 39.5±1.2 | 46.8±1.4 |
| *Astelia alpina* | Asteliaceae | Herb | –13.9±0.7 | –9.8±0.5 | 46.4±1.0 | 60.4±0.8 |
| *Carex breviculmis* | Cyperaceae | Sedge | –12.0±1.2 | –15.3±1.7 | 47.0±0.8 | 59.4±0.7 |
| *Celmisia costiniana* | Asteraceae | Herb | –9.6±0.7 | –7.6±0.7 | 44.9±0.4 | 54.4±0.9 |
| *Epacris paludosa* | Ericaceae | Shrub | –18.2±1.2 | –19.4±1.1 | 45.2±1.7 | 63.0±2.5 |
| *Eucalyptus pauciflora (ssp niphophila)* | Myrtaceae | Tree | –8.7±0.3 | –9.1±0.4 | 39.0±0.7 | 47.7±0.8 |
| *Ewartia nubigena* | Asteraceae | Mat | –13.1±1.8 | –11.9±1.9 | 43.0±0.9 | 56.1±2.2 |
| *Grevillea australis* | Proteaceae | Shrub | –11.3±1.1 | –10.8±1.8 | 44.1±1.4 | 55.4±1.9 |
| *Hovea montana* | Fabaceae | Shrub | –18.5±0.3 | –8.0±2.5 | 44.6±1.3 | 63.1±1.3 |
| *Luzula acutifolia* | Juncaceae | Rush | –8.6±0.4 | –10.3±1.3 | 44.2±2.3 | 52.9±2.3 |
| *Nematolepis ovatifolia* | Rutaceae | Shrub | –7.5±0.4 | –8.1±0.3 | 45.5±3.4 | 53.0±3.7 |
| *Olearia brevipedunculata* | Asteraceae | Shrub | –14.3±1.3 | –11.9±1.7 | 43.6±0.6 | 57.9±1.4 |
| *Orites lancifolius* | Proteaceae | Shrub | –8.6±0.4 | –8.5±0.5 | 45.2±1.7 | 53.8±1.7 |
| *Oxylobium ellipticum* | Fabaceae | Shrub | –16.2±1.6 | –19.4±1.3 | 42.3±1.2 | 58.5±2.4 |
| *Pimelea ligustrina* | Thymelaeaceae | Shrub | –9.6±0.6 | –8.6±0.6 | 37.1±0.4 | 46.6±0.6 |
| *Plantago muelleri* | Plantaginaceae | Herb | –6.0±0.4 | –5.5±0.4 | 39.9±2.9 | 45.9±3.1 |
| *Podocarpus lawrencei* | Podocarpaceae | Shrub | –10.3±0.6 | –13.7±1.7 | 45.8±1.2 | 56.0±1.2 |
| *Prostanthera cuneata* | Lamiaceae | Shrub | –12.5±1.5 | –15.9±1.7 | 39.1±2.0 | 51.7±1.5 |
| *Psychrophila introloba* | Ranuculaceae | Herb | –7.3±0.4 | –6.8±0.5 | 41.6±0.5 | 48.9±0.8 |
| *Dracophyllum continentis* | Ericaceae | Shrub | –13.0±1.4 | –14.4±2.3 | 43.6±0.4 | 56.7±1.3 |
| *Tasmannia xerophila* | Winteraceae | Shrub | –8.4±0.5 | –8.69±0.4 | 44.1±0.7 | 52.4±1.1 |
| *Viola betonicifolia* | Violaceae | Herb | –6.6±1.0 | –7.7±0.7 | 43.5±0.4 | 50.1±1.3 |
| Temperate |  |  |  |  |  |  |
| *Acacia binervata* | Fabaceae | Tree | –12.0±0.7 | –11.8±0.9 | 39.8±1.5 | 51.8±1.0 |
| *Acacia longifolia* | Fabaceae | Tree | –10.2±0.7 | –10.3±0.6 | 43.7±1.7 | 53.8±2.2 |
| *Acmena smithii* | Myrtaceae | Tree | –11.5±0.8 | –6.3±1.6 | 48.0±0.7 | 59.4±1.0 |
| *Backhousia myrtifolia* | Myrtaceae | Tree | –8.5±0.8 | –6.8±1.0 | 48.0±0.4 | 56.5±1.1 |
| *Banksia integrifolia* | Proteaceae | Tree | –6.7±0.5 | –5.1±0.3 | 38.1±1.8 | 44.8±2.2 |
| *Brachychiton acerifolius* | Malvaceae | Tree | –6.6±0.8 | –7.7±0.5 | 48.6±0.4 | 55.2±0.7 |
| *Carex appressa* | Cyperaceae | Sedge | –9.3±0.3 | –5.0±0.7 | 50.1±1.2 | 59.4±1.1 |
| *Cayratia clematidea* | Vitaceae | Climber | –8.0±0.8 | –8.0±0.5 | 46.5±0.2 | 54.6±0.9 |
| *Ceratopetalum apetalum* | Cunoniaceae | Tree | –10.8±1.0 | –8.5±1.8 | 47.3±2.0 | 58.1±3.0 |
| *Cryptocarya microneura* | Lauraceae | Tree | –8.5±1.1 | –11.9±0.3 | 45.4±1.2 | 53.9±1.7 |
| *Doryphora sassafras* | Atherospermataceae | Tree | –8.4±0.7 | –5.1±0.7 | 47.6±0.8 | 55.9±1.1 |
| *Ficus coronata* | Moraceae | tree | –8.7±0.6 | –5.8±0.7 | 46.5±0.5 | 55.2±0.8 |
| *Guioa semiglauca* | Sapindaceae | Tree | –7.7±0.3 | –5.3±0.9 | 50.8±0.4 | 58.5±0.5 |
| *Lomandra longifolia* | Asparagaceae | Sedge | –10.9±1.1 | –6.7±1.2 | 49.5±1.6 | 60.4±2.3 |
| *Luzuriaga cymosum* | Luzuriagaceae | Climber | –10.1±1.7 | –10.9±1.3 | 48.8±0.2 | 58.9±1.7 |
| *Myoporum acuminatum* | Scrophulariaceae | Tree | –9.4±0.6 | –7.3±1.7 | 49.0±0.9 | 58.4±0.9 |
| *Oplismenus hirtellus* | Poaceae | Grass | –4.0±0.7 | –6.2±1.2 | 43.8±1.0 | 47.8±1.0 |
| *Pittosporum undulatum* | Pittosporaceae | Tree | –10.5±1.2 | –8.5±1.4 | 45.2±1.4 | 55.7±2.2 |
| *Podocarpus elatus* | Podocarpaceae | Tree | –8.8±1.1 | –6.6±1.1 | 48.2±0.9 | 57.1±1.9 |
| *Stephania japonica* | Menispermaceae | Climber | –7.1±0.5 | –5.7±1.1 | 48.3±0.6 | 55.4±1.0 |
| *Tasmannia insipida* | Winteraceae | Shrub | –11.7±1.3 | –14.1±0.7 | 48.0±0.6 | 59.7±0.8 |
| *Trochocarpa laurina* | Ericaceae | Shrub | –6.2±0.6 | –3.2±0.5 | 47.9±1.4 | 54.2±1.4 |
| *Viola banksii* | Violaceae | Herb | –7.1±1.6 | –6.0±0.6 | 42.2±0.7 | 49.3±1.6 |
| Desert |  |  |  |  |  |  |
| *Acacia aneura* | Fabaceae | Tree | –13.0±2.2 | –16.2±1.2 | 53.0±4.0 | 66.0±4.0 |
| *Acacia salicina* | Fabaceae | Tree | –13.8±0.8 | –13.6±1.1 | 49.6±0.8 | 63.3±0.9 |
| *Acacia victoriae* | Fabaceae | Tree | –15.6±1.9 | –18.3±0.4 | 45.1±1.8 | 60.7±3.3 |
| *Alectryon oleifolius* | Sapindaceae | Tree | –13.0±0.9 | –13.5±1.1 | 50.3±1.9 | 63.4±2.1 |
| *Atriplex stipitata* | Amaranthaceae | Subshrub | –16.7±0.6 | –20.5±0.4 | 47.2±0.5 | 63.9±1.1 |
| *Brachychiton populneus* | Malvaceae | Tree | –8.8±1.2 | –13.9±1.4 | 47.4±1.9 | 56.2±2.1 |
| *Capparis mitchellii* | Capparaceae | Tree | –13.1±0.6 | –14.7±1.8 | 49.5±1.3 | 62.6±1.5 |
| *Casuarina pauper* | Casuarinaceae | Tree | –14.9±0.3 | –18.0±0.4 | 48.7±0.9 | 63.6±1.3 |
| *Dodonaea viscosa* | Sapindaceae | Shrub | –12.4±1.4 | –14.8±0.7 | 45.4±0.5 | 57.8±1.6 |
| *Eremophila latrobei* | Scrophulariaceae | Shrub | –11.6±0.8 | –18.3±1.1 | 48.5±2.0 | 60.2±2.1 |
| *Eremophila sturtii* | Scrophulariaceae | Shrub | –12.7±1.5 | –14.3±1.4 | 51.3±1.8 | 64.1±2.9 |
| *Eucalyptus coolabah* | Myrtaceae | Tree | –13.3±2.0 | –17.8±0.8 | 46.3±2.4 | 59.6±3.7 |
| *Eucalyptus largiflorens* | Myrtaceae | Tree | –14.3±1.2 | –13.9±0.5 | 45.9±3.1 | 60.2±2.5 |
| *Flindersia maculosa* | Rutaceae | Tree | –13.7±0.7 | –14.9±1.3 | 51.7±4.0 | 65.4±3.6 |
| *Geijera parviflora* | Rutaceae | Tree | –11.3±1.4 | –15.0±1.1 | 51.4±2.2 | 62.8±3.2 |
| *Grevillea striata* | Proteaceae | Shrub | –15.0±1.2 | –17.6±0.3 | 47.6±4.2 | 62.7±4.3 |
| *Myoporum montanum* | Scrophulariaceae | Shrub | –13.9±0.4 | –16.2±1.1 | 48.4±0.9 | 62.2±1.1 |
| *Pimelea microcephala* | Thymelaeaceae | Shrub | –12.2±2.9 | –15.6±0.8 | 48.7±2.1 | 60.9±2.1 |
| *Ptilotus sessilifolius* | Amaranthaceae | Subshrub | –12.7±3.4 | –17.2±2.2 | 51.8±0.5 | 64.6±3.6 |
| *Rhagodia spinescens* | Amaranthaceae | Subshrub | –15.4±2.2 | –18.1±1.2 | 54.9±3.4 | 70.3±4.5 |
| *Sclerolaena birchii* | Amaranthaceae | Subshrub | –13.9±1.5 | –15.4±0.7 | 48.6±1.6 | 62.5±2.8 |
| *Senna artemisioides* | Fabaceae | Shrub | –13.4±2.2 | –18.1±0.6 | 51.9±1.0 | 65.3±2.7 |
| *Sida fibulifera* | Malvaceae | Subshrub | –13.1±2.1 | –15.2±0.9 | 50.5±0.9 | 63.6±2.0 |
| *Solanum esuriale* | Solanaceae | Herb | –11.9±0.9 | –13.5±1.5 | 49.3±1.1 | 61.2±0.8 |

**Table S2.** Local climate variables extracted for species locations across all biomes (see associated spreadsheet).

**Table S3.** Tukey’s Honest Significant Differences pairwise biome contrasts for each thermal tolerance threshold, corresponding to the post-hoc tests should with letters on Figure 2.

| **Cold tolerance threshold: T_crit-cold_ (°C)** | | | | |
| --- | --- | --- | --- | --- |
| **Contrast** | **Difference** | **Lower 95% CI** | **Upper 95% CI** | ***p*-value** |
| Temperate-Alpine | 2.140 | 1.057 | 3.224 | **< 0.001** |
| Desert-Alpine | –2.305 | –3.360 | –1.250 | **< 0.001** |
| Desert-Temperate | 2.748 | 1.394 | 4.101 | **< 0.001** |
| **Freezing tolerance: T_nucleation_ (°C)** | | | | |
| **Contrast** | **Difference** | **Lower 95% CI** | **Upper 95% CI** | ***p*-value** |
| Temperate-Alpine | 3.212 | 2.046 | 4.379 | **< 0.001** |
| Desert-Alpine | –5.300 | –6.478 | –4.122 | **< 0.001** |
| Desert-Temperate | –8.512 | –9.719 | –7.306 | **< 0.001** |
| **Heat tolerance threshold: T_crit-hot_ (°C)** | | | | |
| **Contrast** | **Difference** | **Lower 95% CI** | **Upper 95% CI** | ***p*-value** |
| Temperate-Alpine | 3.447 | 2.108 | 4.785 | **< 0.001** |
| Desert-Alpine | 6.194 | 4.891 | 7.498 | **< 0.001** |
| Desert-Temperate | 2.748 | 1.394 | 4.101 | **< 0.001** |
| **Thermal Tolerance Breadth: TTB (°C)** | | | | |
| **Contrast** | **Difference** | **Lower 95% CI** | **Upper 95% CI** | ***p*-value** |
| Temperate-Alpine | 1.370 | –0.4146 | 3.155 | 0.169 |
| Desert-Alpine | 8.563 | 6.825 | 10.301 | **< 0.001** |
| Desert-Temperate | 7.193 | 5.389 | 8.997 | **< 0.001** |

**Table S4.** Results of a linear mixed-effects regression model that tested whether T_nucleation_ was different to T_crit-cold_ in each biome.

|  | **T_nucleation_ – T_crit-cold_ (°C)** | | | | | | |
| --- | --- | --- | --- | --- | --- | --- | --- |
| **Fixed** | **Coefficient ± SE** | ***t-*value** | ***p-*value** | **Random** | **Coefficient** | ***R^2^* type** ^*^ | ***R^2^* value** |
| (Intercept,  Biome: alpine) | 0.136 ± 0.491 | 0.277 | 0.782 | Growth form | 0.000 | *R*^2^*m* | 0.155 |
| Biome (temperate) | 1.134 ± 0.698 | 1.626 | 0.105 | Species | 1.774 | *R*^2^*c* | 0.328 |
| Biome (desert) | –2.932 ± 0.697 | –4.205 | **<0.001** | Phylogeny | 0.000 | *R*^2^*c* - *R*^2^*m* | 0.173 |
|  |  |  |  | Residual | 3.500 |  |  |

^*^*R*^2^*m* represents the variance explained by all fixed effects, *R*^2^*c* represents the variance explained by all fixed and random effects, and *R*^2^*c* – *R*^2^*m* represents the variance explained by all random effects.

**Table S5.** Loadings and variance explained by the first and second Principal Components (PC1 and PC2) that explain 99.6% of the variance among local climate variables. Bold indicates dominant variables.

|  | **Loadings** | |
| --- | --- | --- |
| **Local climate variable** | **PC1** | **PC2** |
| MAP | **–0.498** | –0.158 |
| MinT | 0.382 | **–0.624** |
| MaxT | **0.506** | 0.098 |
| Trange | 0.334 | **0.713** |
| MAT | **0.489** | –0.261 |
| Variance explained | 77.2% | 22.4% |
| Cumulative variance | 77.2% | 99.6% |

**Table S6.** Results of tests for spatial autocorrelation within and among biomes across the thermal tolerance thresholds using Moran’s *I*.

|  | **All biomes** | | **Alpine** | | **Temperate** | | **Desert** | |
| --- | --- | --- | --- | --- | --- | --- | --- | --- |
| **Trait** | **Moran’s *I*** | ***p-*value** | **Moran’s *I*** | ***p-*value** | **Moran’s *I*** | ***p-*value** | **Moran’s *I*** | ***p-*value** |
| T_crit-cold_ | –0.0098 | 0.679 | –0.0062 | 0.960 | –0.0330 | 0.349 | 0.0271 | 0.268 |
| T_nucleation_ | 0.0104 | 0.472 | –0.0010 | 0.800 | –0.0120 | 0.904 | 0.0039 | 0.718 |
| T_crit-hot_ | –0.0085 | 0.738 | 0.0068 | 0.585 | 0.0407 | 0.053 | 0.0204 | 0.367 |
| TTB | 0.0024 | 0.769 | 0.0010 | 0.743 | 0.0270 | 0.160 | –0.0245 | 0.604 |

**Table S7.** Model AIC and weights for each physiological thermal tolerance threshold and TTB.

| **Trait** | **Model fixed effect structure** | ***npar*** | **AIC** | **Weight** |
| --- | --- | --- | --- | --- |
| T_crit-cold_ | PC1 + PC2 | 7 | 1928.0 | 0.049 |
|  | PC1 + PC1^2^ + PC2 | 8 | 1923.3 | **0.524** |
|  | PC1 + PC2 + PC2^2^ | 8 | 1925.4 | 0.180 |
|  | PC1 + PC1^2^ + PC2 + PC2^2^ | 9 | 1924.8 | 0.248 |
| T_nucleation_ | PC1 + PC2 | 7 | 1805.0 | 0.076 |
|  | PC1 + PC1^2^ + PC2 | 8 | 1803.8 | 0.136 |
|  | PC1 + PC2 + PC2^2^ | 8 | 1800.9 | **0.566** |
|  | PC1 + PC1^2^ + PC2 + PC2^2^ | 9 | 1802.8 | 0.223 |
| T_crit-hot_ | PC1 + PC2 | 7 | 2126.6 | **0.525** |
|  | PC1 + PC1^2^ + PC2 | 8 | 2128.5 | 0.203 |
|  | PC1 + PC2 + PC2^2^ | 8 | 2128.6 | 0.194 |
|  | PC1 + PC1^2^ + PC2 + PC2^2^ | 9 | 2130.4 | 0.078 |
| TTB | PC1 + PC2 | 7 | 2307.4 | 0.192 |
|  | PC1 + PC1^2^ + PC2 | 8 | 2305.6 | **0.480** |
|  | PC1 + PC2 + PC2^2^ | 8 | 2307.9 | 0.151 |
|  | PC1 + PC1^2^ + PC2 + PC2^2^ | 9 | 2307.6 | 0.177 |

**Table S8.** Results of tests for phylogenetic signal in physiological thermal tolerance thresholds.

| **Trait** | ***C*_mean_** | ***p-*value** | ***I*** | ***p-*value** | ***K*** | ***p-*value** | **λ** | ***p-*value** |
| --- | --- | --- | --- | --- | --- | --- | --- | --- |
| T_crit-cold_ | 0.134 | 0.052 | 0.006 | 0.243 | 0.112 | 0.342 | 0.516 | 0.433 |
| T_nucleation_ | 0.104 | 0.085 | –0.004 | 0.355 | 0.103 | 0.423 | 0.314 | 1.000 |
| T_crit-hot_ | –0.182 | 0.992 | –0.070 | 0.958 | 0.045 | 0.968 | 0.000 | 1.000 |
| TTB | –0.075 | 0.757 | –0.011 | 0.425 | 0.113 | 0.362 | 0.000 | 1.000 |
